# Molecular signatures of hyperexcitability and lithium responsiveness in bipolar disorder patient neurons provide alternative therapeutic strategies

**DOI:** 10.1101/2023.07.21.550088

**Authors:** Anouar Khayachi, Malak Abuzgaya, Yumin Liu, Chuan Jiao, Kurt Dejgaard, Lenka Schorova, Anusha Kamesh, Qin He, Yuting Cousineau, Alessia Pietrantonio, Nargess Farhangdoost, Charles-Etienne Castonguay, Boris Chaumette, Martin Alda, Guy A. Rouleau, Austen J. Milnerwood

**Author notes:** Equal contribution (Malak Abuzgaya, Yumin Liu).

## Abstract

Bipolar disorder (BD) is a multifactorial psychiatric illness affecting about 1% of the world population. The first line treatment, lithium (Li), is effective in only a subset of patients and its mechanism of action remains largely elusive. In the present study, we used iPSC-derived neurons from BD patients responsive (LR) or not (LNR) to lithium and combined electrophysiology, calcium imaging, biochemistry, transcriptomics, and phosphoproteomics to report mechanistic insights into neuronal hyperactivity in BD, and Li’s mode of action. We show a selective rescue of neuronal hyperactivity by Li in BD LR neurons through changes in Na^+^ currents. The whole transcriptome sequencing revealed altered gene expression in BD neurons in pathways related to glutamatergic transmission, and Li selectively altered those involved in cell signaling and ion transport/channel activity. We found the therapeutic effect of Li in BD LR patients was associated with Akt signaling and confirmed that an Akt activator mimics Li effect in BD LR neurons. Further, we showed that AMP-activated protein kinase (AMPK) reduces neural network activity and sodium currents in BD LNR patients. These findings suggest the potential for novel treatment strategies in BD, such as Akt activators in BD LR cases, and the use of AMPK activators for BD LNR patients.

## INTRODUCTION

Bipolar Disorder^1^ (BD) is a major psychiatric illness affecting 1-3% of the global population^2^. It is ranked in the top 10 causes of disability and mortality worldwide, largely due to a 15% suicide rate, in addition to cardiovascular and metabolic comorbidities^3–5^. Typically, BD has life-long presentation characterized by alternations of mania and depression, often associated with psychosis and self-harm^1^. Its etiology is likely multifactorial involving both genetic and environmental factors. Previous studies show BD to be a highly heritable disease (up to 70%), and successful treatment options often work within affected families, suggesting a strong genetic component^6–8^. Genome Wide Association Studies (GWAS) have revealed many single nucleotide polymorphisms (SNPs) as risk factors for BD, including genes encoding proteins of synaptic, calcium, and second messenger signaling pathways^9^. Despite these advances, the links between familial presentation, GWAS hits, neurobiological mechanisms, symptoms, and successful treatments, remain unresolved. The unknown cause and genetic heterogeneity of BD are major challenges to research efforts, especially those aimed at producing valid disease models in which to explore underlying mechanisms and potential new therapeutics.

Clinical management is also challenging, due to heterogeneous symptom presentation, phenocopies and similarity to other disorders (e.g. unipolar depression and schizophrenia). Care is further complicated by frequent comorbidity with anxiety or substance abuse among others, that often lead to misdiagnosis and incorrect treatment at early stages. Long-term use of mood stabilizers is the cornerstone of BD clinical management, but clinicians often rely on a “trial-and-error” strategy to find an effective treatment. Consequently, it can take many months or even years to achieve mood stabilization. Despite a low therapeutic index, the mood stabilizer lithium (Li) remains the gold standard treatment for prevention of the illness episodes; it also effectively reduces suicide and overall mortality rates^10^ in the ∼30% of BD patients who respond to it. Aside Li, other medications have come from drug repurposing (e.g. anticonvulsants valproate and lamotrigine), which highlights our poor understanding of BD pathophysiology, and the struggle to develop novel therapies through mechanistic insights.

To better understand Li as a treatment, and to potentially extrapolate its effects to other compounds with a larger therapeutic index, two major questions remain; i) how is Li effective, and ii) why so for only some patients^11, 12^. Li’s mode of action is unknown, with multiple direct and indirect targets and substantial cellular effects^13^. Li variously affects several kinase / phosphatase cascades e.g., it inhibits protein kinase C (PKC) and glycogen synthase 3b (GSK3b)^14–17^, activates protein kinase B (Akt/PKB), and inhibits inositolpoly-(IMPase) and inositol mono-phosphatase (IMPase)^13^. These molecular changes likely alter several intracellular signaling cascades that modulate neuronal excitability, synaptic transmission, gene expression, and chronobiology. In mouse primary cortical neuron cultures, we previously found long-term Li exposure decreased neuronal sodium (Na^+^) conductance, action potential firing, and calcium (Ca^2+^) flux in individual neurones. Furthermore, Li exposure had effects on the neural network, altering downstream Ca^2+^ signal cascades, decreasing excitatory and increasing inhibitory synapse number and activity^14^.

Induced pluripotent stem cell (iPSC) technology facilitates personalised study of neuropsychiatric disorders at the cellular and molecular level, with a non-invasive opportunity to study the function of neuronal cells derived from individual patients. Moreover, data obtained can be aligned with known genetic profiles and associated clinical data e.g., lithium responsiveness. Our previous and others’ recent reports^18, 19^ found hyperexcitability in iPSC-derived neurons from BD patients, which was Li selectively reversed in cells from clinically responsive patients. Li effects were associated with changes to Na^+^ and potassium (K^+^) currents, but the mechanisms underlying neuronal hyperactivity and Li’s mode of action were unclear.

Here, we used iPSC-derived forebrain neurons from healthy individuals (Ctl), Li-responsive (LR) and Li-non-responsive (LNR) BD patients. We combined electrophysiology, calcium imaging, biochemistry, transcriptomics, and phosphoproteomics to provide mechanistic insights into neuronal hyperactivity in BD, and Li’s mode of action in BD LR neurons. We aim to guide the search for biomarkers of BD, predictors of Li responsiveness, and potential novel therapy for BD LNR patients. We confirm neuronal hyperactivity in BD neurons and selective rescue by Li in BD neurons from LR (and not BD LNR) patients, through changes in Na^+^ currents. Whole transcriptome sequencing revealed altered gene expression in BD neurons in pathways related to glutamatergic synaptic activity, and Li selectively altered those involved in cell signaling and ion transport/channel activity. We found the therapeutic effect of Li in BD LR patients was associated with Akt signaling and confirmed that an Akt activator mimics Li effect in BD LR neurones. Further, we found that AMP-activated protein kinase (AMPK) and reduces neural network activity and sodium currents in BD LNR patients. Here, we propose the potential for novel treatment strategies in BD, such as Akt activators to mimic the therapeutic effect of Li with less risk of toxicity in BD LR cases, and the use of AMPK activators for BD LNR patients.

## RESULTS

### The composition of neuron cultures of Li-responsive and -non-responsive BD patients is similar to those of controls

We sourced blood from BD patients in well-characterized Li-responding and Li-non-responsive families, and age- and sex matched healthy individuals as controls (Ctl; no neuropsychiatric disease at the time of sample collection). These samples were used to isolate lymphocytes or peripheral blood mononuclear cells from which iPSCs were produced. Samples were selected based on clinical data according to the Alda scale for Li-responsiveness^20, 21^ with a score of 9 to 10/10 for BD Li-responsive (BD LR) and 0 to 3/10 for BD Li-non-responsive (BD LNR) patients (*table 1*). All iPSCs lines were differentiated into neural progenitor cells (NPCs) with a minimum of ∼90% of Nestin and Sox2+ve cells (*extended Fig. 1A-B*) selected for subsequent neuronal differentiation. At 4 weeks post-differentiation, neuronal phenotypes were confirmed by immunostaining for the neuron-specific marker beta-III tubulin/Tuj1, which demonstrated ∼75% of all cells were Tuj1+ve / neuronal (*Fig. 1A*). Approximately 65% of all cells were positive for both Tuj1 and the vesicular glutamate transporter 1(Vglut1), indicating that majority of the cells were neurones, and these neurones were predominantly glutamatergic (*Fig. 1B*); There were no differences in the cellular composition of Ctl and BD cultures.

**Figure 1:**
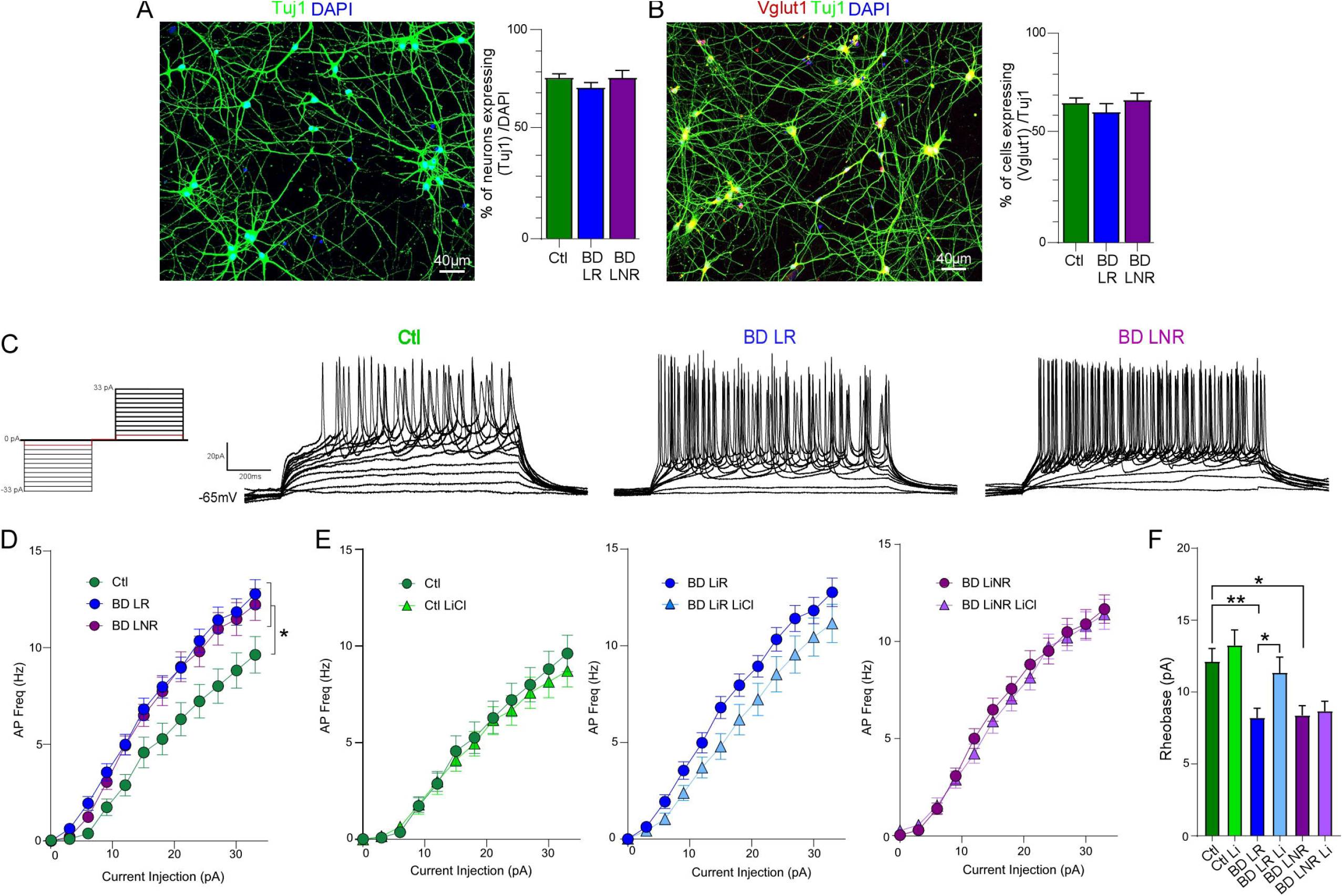
Hyperexcitability in iPSC-derived neurons from BD patients is selectively reversed by Li treatment in Li-responsive patient cell lines. ***A-B)*** Characterization of iPSC-derived neuron cultures at 4 weeks post-neuronal differentiation. ***A)*** cultures were stained for the neuronal marker (Tuj1) and nuclei (DAPI), in all lines. ***B)*** Glutamatergic neurons were identified by positive staining for the vesicular glutamate transporter (VGluT1). ***C)*** Example of current-clamp membrane potential responses to 3pA increasing positive and negative current injection steps from 0 to 33pA (left) in Ctl, BD LR & BL LNR neurons ***D-E)*** Quantification of AP frequency (spikes/s) *vs* positive current injection, and rheobase. ***D)*** BD (LR & LNR) neurons exhibit higher AP frequencies than Ctl neurons at all current steps above rheobase. ***E)*** Ctl, BD LR and LNR neurons +/-Li (1.5mM; 7d). Li had no effect on Ctl AP frequency plots, but reduced AP frequency in neurones from BD LR patient cell lines. No differences in AP frequency plots were produced by Li in neurons from BD LNR patient lines. ***F)*** Mean neuronal rheobase was significantly lower in neurons from BD LR & LNR patient lines, relative to Ctl lines; Li significantly increased rheobase in BD LR neurons, but not BD LNR (or Ctl) neurons. Data is percent +ve cells (mean ±SEM of each group, sampled from ∼40 images fields in each of 4-5 lines per group) for A-B, and mean ±SEM of ∼50 neurons per condition from 4-5 different lines per group at 28 to 32 days post-neuronal differentiation *(**D-F**)*. Statistical comparisons were by one-way *(**A, B&F**)* or two-way ANOVA (***D-E***) with Bonferroni’s multiple comparison post-test; \**p*< 0.05.

**TABLE 1:**
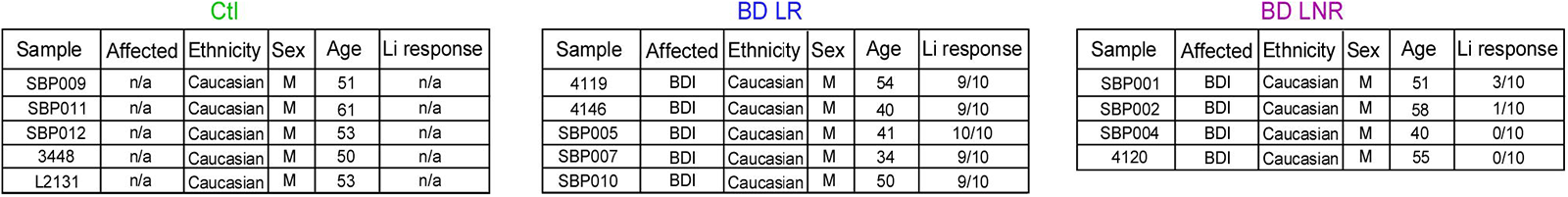
Origin of patient samples from BD patients, clinical response to Li treatment, and matched controls. Sample ID: all samples are sourced from blood SBP001, 2, 4, 5, 7, 9, 10, 11, 12 (iPSC are derived from lymphocytes) and 3448, L2131, 4119, 4146, 4120 (iPSC are derived from peripheral blood mononuclear cells); n/a = not applicable; BDI = Bipolar Disorder type I; M = Male; Age = years; Li response as rated by clinical assessment with the standardized Alda scale.

### BD patient neurons are hyperexcitable, and Li specifically reduces hyperexcitability in Li-responsive patient neurons

To assess the physiology of human neurones, we conducted whole-cell current-clamp recordings of neuronal intrinsic membrane properties and action potential (AP) characteristics. Current/Voltage (I/V) plot shows that membrane capacitance was significantly different in BD vs Ctl neurons and upon Li treatment exclusively in BD LR neurons (see supplemental extended Fig. 1 I-J). Upon positive current injection, AP firing frequency was increased, and rheobase (amount of current needed to trigger an AP) was significantly reduced, in neurons from BD patients relative to controls (*Fig. 1C, D, E & F*), confirming the previous observation of hyperexcitability in both LR and LNR BD neurones^18, 19^. Chronic application of Li (1.5mM for 7 days) resulted in significantly fewer APs for treatment effect in BD LR neurons and increased rheobase, specifically in BD neurones from LR patients; Li had no effect in Ctl or BD LNR neurons (*Fig. 1E, F and extended Fig1.D, E*).

In a subset of the same cells, we also conducted whole-cell voltage-clamp recording of spontaneous glutamatergic (holding voltage (Vh) -60mV) and GABAergic (Vh at 0mV) postsynaptic currents (sEPSCs and sIPSCs) respectively. We found a significant increase in sEPSC frequency in BD LR and LNR neurons, which was selectively reversed by Li treatment (1.5mM, 7 days) in BD-LR neurons, but found no change to sIPSC frequency (extended Fig. 2A-*D*).

Together, the data demonstrate intra-neuronal and excitatory network hyperactivity that is specifically reduced in only BD LR neurons by Li.

### Increased Na^+^ current in neurons from BD LR patients is rescued by Li

To further understand hyperexcitability in BD neurons we assayed Na^+^ and K^+^ currents by whole-cell voltage clamp (*Fig. 2A*). We found a significant increase in Na^+^ current input/output curves, specifically in BD LR neurons, in concert with increased peak Na+ conductance that trended to significance (*Fig. 2B&C*). There was a weaker trend to increased slow and fast K+ current flux in both BD LR and BD LNR, relative to controls (*Fig. 2B-E*). Li application resulted in a marked, significant, reduction of Na^+^ current input/output curves and peak conductance in BD LR neurons (*Fig. 2F-G*) but had no effect in BD LNR or Ctl neurons (*extended Fig. 3B-E*). To ensure consistent Li effects within each patient line, we compare each separately and confirmed a significant reduction of Na^+^ current in neurons from each BD LR patient cell line *(extended Fig. 3A);* whereas no reductions were observed in neurones from BD LNR or Ctl cell lines *(Fig. 3C, E)*. While trends to altered fast and slow K^+^ currents in BD neurones did not reach significance, separate quantification of Li effects in each patient line showed a consistent and significant reduction of fast and slow K^+^ current exclusively in BD LR neurons (*extended Fig. 3G, I, K, M, O, Q)*. The data demonstrate Na^+^ channel overactivation in BD LR neurons, and that Li treatment specifically reduces Na^+^ and K^+^ channel conductance in BD LR neurons.

**Figure 2:**
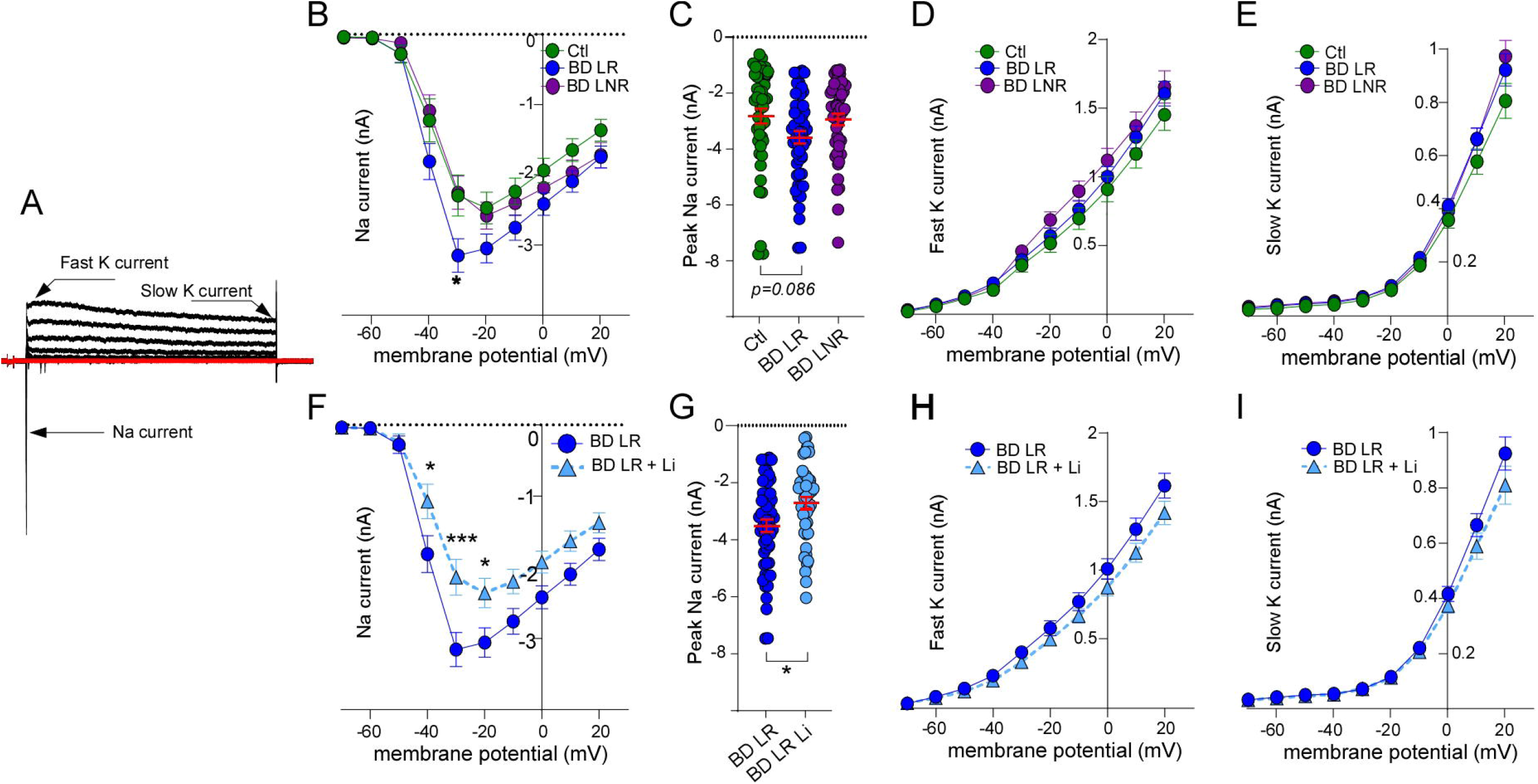
Increased sodium current in neurons from BD LR patient lines is normalised by Li. ***A)*** Example of voltage-clamp Na+ and K+ currents in response to 10mV increasing positive voltage steps from -70mV to 20mV in Ctl neuron. Arrows indicate Na+ current and peak amplitude of fast and slow K+ currents. Quantification of voltage-dependence of ***B)*** peak sodium current *vs* V*m **C)*** maximum Na+ peak, ***D)*** fast and ***E)*** slow potassium current *vs* V*m* in Ctl, BD LR and BD LNR patient line neurons. ***F&G)*** Li (1.5mM, 7d) reduced Na+ current peaks in BD LR neurons. ***H)*** Mean fast potassium and ***I)*** slow potassium current x voltage plots in BD LR neurons were reduced. Data are means±SEM of ∼50 neurons per condition from 4-5 different lines per group at 28 to 32 days post-neuronal differentiation. Statistical comparisons were by two-way (**B**, **D**, **E**, **F**, **H**, **I**) or one-way ANOVA (**C-G**) with Bonferroni’s multiple comparison post-test, and Welch’s t test was (**D**) ***p< 0.001, *p< 0.05.

**Figure 3:**
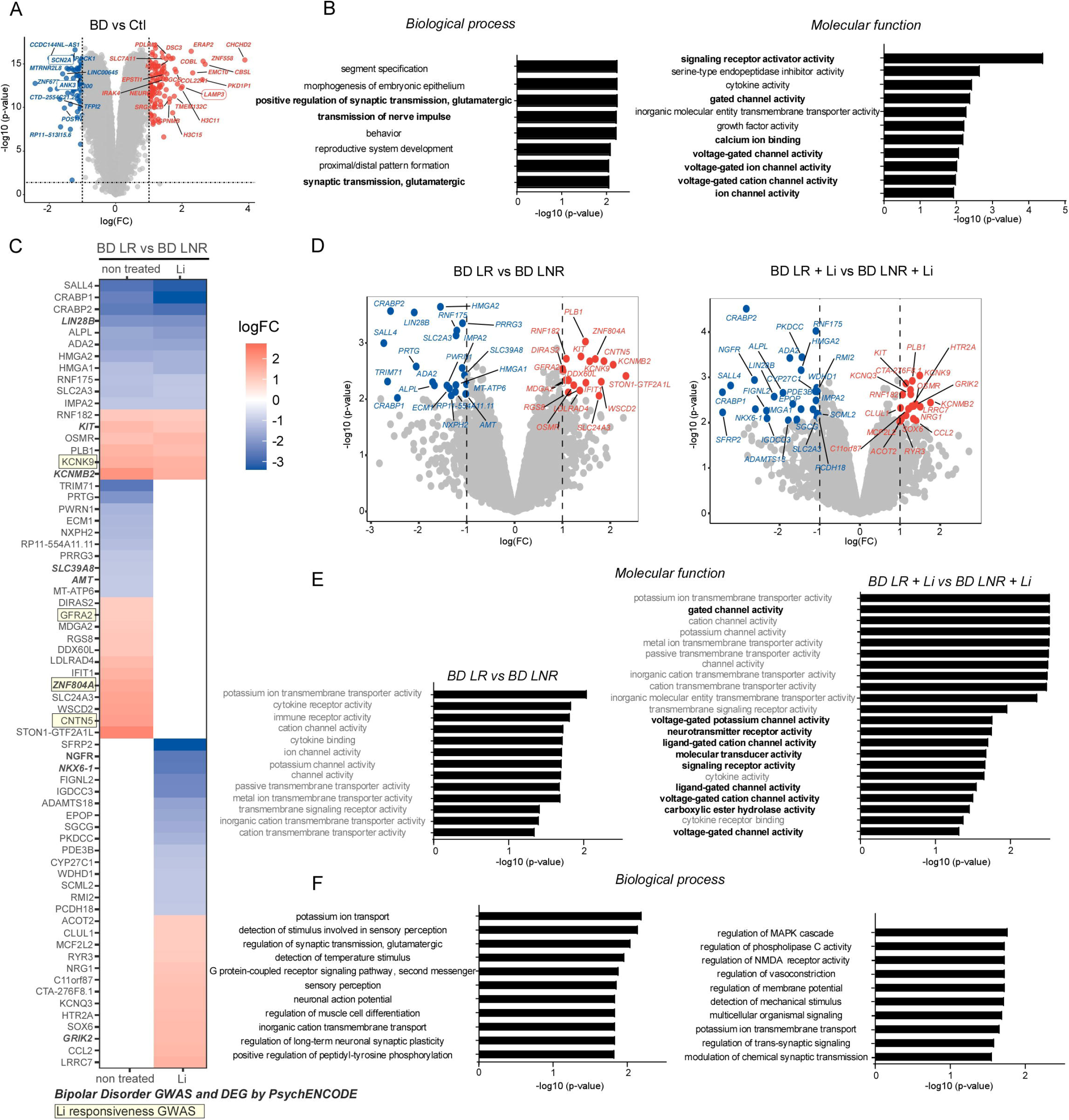
Differential gene expression in neurons from BD patients and identification of gene expression changes by Li application in LR vs LNR patients. ***A)*** Volcano plot shows gene expression in neurons from BD patients *vs.* those from Ctl at ∼30 days post-neuronal differentiation (data from 5 individual lines for Ctl and all 9 BD lines LR & LNR). Differentially expressed genes (DEGs) were identified as those with a p<0.05 and log p-value >2 with log FC >1. Framed genes are those previously identified by BD GWAS (see result section). ***B)*** Biological processes and molecular functions of DEG in BD neurons grouped by GO analysis. ***C&D)*** Heatmap (C) and volcano plots (D) of DEGs in BD LR and BD LNR neurons, and non- and Li-treated conditions revealed an overlap of 17 genes between -/+ Li conditions, 22 DEGs were no longer detectable, and 28 DEGs were newly detected in Li treated neurons. Highlighted and framed genes are those identified in previous BD and Li responsiveness GWAS or RNAseq studies (see result section). **E)** Molecular functions of DEGs identified between LR & BD LNR neurons (left) and -/+ Li (right) found molecular functions that were not (in gray) or were affected (in black bold) by Li. ***F)*** Biological processes of genes altered by Li in BD LR neurons. Data from 4-5 lines per group/condition and DEGs identified as those with p<0.05 and log p-value >2 with log FC >1, as in A.

### Differential expression of ion transport & synaptic regulatory genes in BD neurons, and specific effects of Li on gene expression in BD LR & LNR patient neurons

To identify cellular processes underlying hyperexcitability in BD neurons, and effects of Li that are general or specific to BD LR neurons, we performed whole transcriptome sequencing from all patient lines. We compared neurons of Ctl *vs*. BD, BD LR *vs*. BD LNR, and BD LR *vs*. BD LNR -/+ Li.

We identified 177 differential expressed genes (DEGs) in BD, relative to Ctl, in which 107 were upregulated and 70 downregulated. Identified genes were noticeably involved in neuronal excitability, regulation of glutamatergic synapse function, neurotransmitter receptor and Ca^2+^ signaling, and ion transport / channel activity. Hits included GRIN2B, KCNJ6 and CACNG8 that encode for glutamate ionotropic receptor NMDA subunit 2B, K^+^ inwardly rectifying channel subfamily J member 6, and Ca^2+^ voltage gated channel auxiliary subunit 8, respectively. Of note, previously some identified hits in BD GWAS <u>GWAS Catalog</u> <u>(ebi.ac.uk)</u> were identified here as highly differentially expressed in BD neurons (*Fig. 3A*): SCN2A, ANK3, and LAMP3 which encode Na^+^ voltage gated channel alpha subunit 2, Ankyrin 3, and lysosome associated protein 3.

Accordingly, gene ontology (GO) analysis for biological processes highlighted neuronal-specific transcripts, including those for glutamatergic transmission, as significantly altered in BD neurons, relative to Ctl. Similarly GO molecular function analysis suggested dysregulation of neurotransmitter receptor and Ca^2+^ signaling, in addition to ion channel activity (*Fig. 3B*). Comparison of BD LR with BD LNR, and BD LR with BD LNR -/+ Li conditions identified an overlap of 17 genes between -/+ Li (i.e. no Li effect). Then, 22 DEGs that were no longer observed in Li condition (i.e. effect reversed by Li) and finally, 28 DEGs that were solely found with Li treatment (*Fig. 3C&D*). The genes most significantly altered by Li are involved in neuronal excitability, regulation of glutamatergic synaptic transmission, K^+^ transport, G-protein coupled receptor signaling pathway and Ca^2+^ signaling. For example, SLC24A3 (solute carrier family 24 member 3) encoding the plasma membrane Na^+^/Ca^2+^ exchanger, KCNQ3 (potassium voltage-gated channel), and GRIK2 (glutamate ionotropic receptor kainite type subunit 2) all regulate neuronal excitability (*Fig. 3E, F)*.

Interestingly, several genes changes we found to be reversed by Li treatment in BD LR neurons have been previously identified in BD or Li response GWAS <u>GWAS Catalog</u> <u>(ebi.ac.uk)</u>. Of note, SLC39A8 and AMT were identified as hits in a BD GWAS, GFRA2 and CNTN5 were identified as hits in a GWAS of Li responsiveness, whereas ZNF804A (encoding Zinc Finger Protein 804A) was identified in both GWAS studies, and as a DEG here (*Fig. 3C*). It is important to note that RNA levels do not correlate directly with protein levels or functional state. While DEGs highlight changes to potentially important processes, they do not inform on the molecular and biochemical status of proteins that might underlie molecular and physiological differences in BD, or the cell biological response to Li. That said, clear differences in the expression of genes related to neuronal and synaptic transmission were identified between Ctl and BD neurons, and the data suggest Li acts on intracellular signal transduction and second messenger systems that modulate gene expression. Thus, gene expression may be secondarily, rather directly, altered by the presence of Li (*Fig. 3E, F)*.

### Li promotes Akt signaling pathway activation in neurons from BD LR patients

The activity of many kinases has been implicated in BD and Li effects^13^, suggesting the phosphorylation state that governs both their activity, and that of their substrates, should form a consistent pattern. We therefore examined the phosphorylation profile of proteins to elucidate the signaling pathways altered in BD neurons, and which pathways respond to Li using unbiased label-free quantitative phosphoproteomics with nanoflow liquid chromatography tandem mass spectrometry (LC-MS/MS). Phosphopeptides, were enriched by titanium dioxide (TiO2) in neuronal lysates from 4 patient lines per group to compare Ctl *vs.* BD, BD LR *vs.* BD LNR and BD LR *vs.* BD LNR -/+ Li.

We identified 6478, 7355 and 8559 phosphopeptides in Ctl, BD LR and BD LNR neurons, respectively *(extended Fig. 4A),* belonging to ∼4000 identified phosphoproteins. No change was observed in the overall percentage of phosphorylated serine, threonine, and tyrosine sites between groups, demonstrating no drastic change to global phosphorylation levels; ∼85% of serine, ∼14% of threonine, and ∼1% of tyrosine residues were phosphorylated in all samples of Ctl, BD LR and BD LNR -/+ Li *(extended Fig. 4B)*. This finding is also a positive indicator of consistent sample preparation and quality control.

Statistical analyses identified 36, 15, and 43 proteins that were differentially phosphorylated between Ctl vs BD LR, Ctl vs BD LNR, and BD LR vs BD LNR, respectively *(Fig. 4A)*. Using a combination of bioinformatic resources and available proteomics database (phosphoSitePlus^22^, Phospho.ELM^23^, Networkin^24^ and ‘KEA substrates of kinase’ in Harmonizome^25^) we determined which kinases are believed to phosphorylate the identified substrates, and group them to the pathways involved. We identified 5 main kinase signaling cascades: mitogen-activated protein kinase (MAPK), protein kinase C (PKC), glycogen synthase 3 (GSK3), protein kinase A (PKA) & AMP-activated protein kinase (AMPK) (both latter are strongly associated with intracellular adenosine monophosphate (AMP) levels) and protein kinase B (Akt) activity. Of the 27 proteins in the BD LR group with upregulated phosphorylation, we found 44% related to MAPK signaling, and 17%, 14%, 14% and 11% related to PKC, GSK3, PKA&AMPK and Akt signaling, respectively *(Fig. 4A)*. Enrichment analysis (by g:profiler^26^) also suggested an upregulation of small GTPase binding proteins in BD patient neurons, which are similarly involved in the regulation of intracellular signal transduction and insulin signaling *(Fig. 4B)*. Interestingly, accumulating evidence suggest that disrupted insulin signaling is involved in bipolar disorder (BD) pathogenesis and/or mood stabilizer effects^5, 27, 28^.

**Figure 4:**
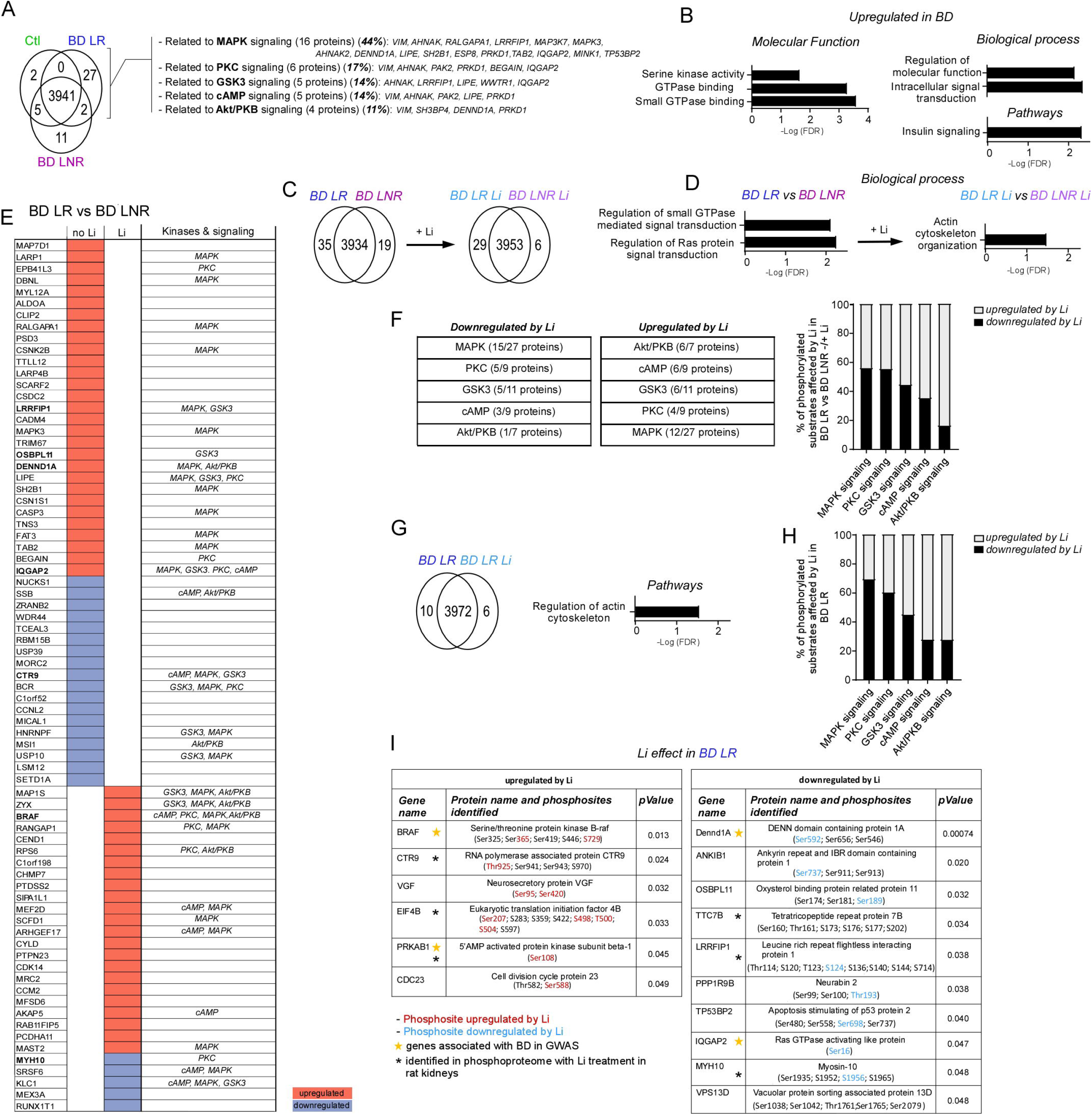
Phosphoproteomic signature of BD and specific effects of Li in BD LR patient neurons. ***A)*** Venn diagram of phosphoproteins identified specific and common to Ctl, BD LR, and BD LNR neurons (left) and grouping of the BD-specific phosphoproteins listed by summary of their associated kinase / signaling cascades. ***B)*** Phosphoproteins unregulated in BD (*vs.* Ctl) grouped by molecular function and biological processes. ***C)*** Venn diagram of phosphoproteins common or specific to BD LR and BD LNR patient neurons in non-treated and Li conditions. ***D)*** Biological processes implicated in BD LR vs BD LNR and by Li. ***E)*** Heat map shows differentially detected phosphoproteins in BD LR vs BD LNR neurons between non- and Li-treated conditions. 47 differentially detected phosphoproteins are no longer present, and 28 phosphoproteins are newly detected, in Li treated BD LR vs BD LNR neurons. Known kinases and signaling cascades for each identified phosphoprotein (substrate). Phosphoproteins linked to kinases also identified in BD LR -/+ Li conditions are shown in bold. ***F)*** Table and summary graph showing the number and percentage of identified substrates (in E) that are phosphorylated by MAPK, PKC, GSK3, AMP and Akt signaling, and which is up or downregulated by Li treatment. ***G)*** Venn diagram showing differentially detected phosphoproteins in BD LR -/+ Li, the pathways implicated, and ***H)*** the percentage of substrates regulated by MAPK, PKC, GSK3, AMP and Akt in non-vs Li-treated cells. ***I)*** Table listing the differentially detected phosphosites in the most significantly altered substrates identified from BD LR neurons -/+ Li. Phosphosites highlighted in red or blue represent upregulated or downregulated phosphorylation by Li, respectively. Yellow star indicates association with BD in previous GWAS studies with “*GWASdb SNP-Disease Associations*” in harmonizome; black star highlights proteins regulated by Li identified in previous phosphoproteomics study (ref: https://esbl.nhlbi.nih.gov/Databases/Lithium-Phospho-IMCD/). All data are from 4 lines per group/condition at ∼30 days post-neuronal differentiation. Comparisons of >2 groups were conducted by one-way ANOVA, and separately by t-test for direct comparisons. Significance set at p< 0.05.

To isolate Li effects from beneficial neuronal (and clinical) Li-responsiveness, we compared BD LR *vs.* BD LNR -/+ Li, and BD LR *vs.* BD LR Li. We found that similarly to BD itself, Li affects small G-protein function and actin cytoskeleton organization. We identified 54 and 35 differentially phosphorylated proteins in BD LR *vs.* BD LNR and BD LR Li *vs.* BD LNR Li, respectively. There was an overlap of 7 proteins that were differentially phosphorylated in BD regardless of Li treatment (i.e. no effect of Li on the phosphorylation; *extended Fig. 4D*). Then, 47 proteins were no longer detected as different in Li (i.e. phosphorylation differences reversed by Li; 18 upregulated and 29 downregulated by Li). Finally, 28 proteins that were only differentially phosphorylated in Li condition (i.e. phosphorylation altered by Li; 5 downregulated and 23 upregulated by Li; *Fig. 4E)*. Linking the kinases that differentially phosphorylate the identified BD substrates reveal which signaling pathways are modulated by Li in BD LR patient neurons, and to what extent.

Each kinase/signaling cascaded we identified contains substrates that are downregulated and upregulated by Li, which seems to be paradoxical. Substrate phosphorylation is not only dictated by kinase activity, but also by many other determinants including other lower affinity kinases, cofactors, and subcellular localization / membrane associations. That said, the ratio between downregulation and upregulation of all substrates of a kinase or signaling pathway generally indicates the direction in which Li is altering the signaling pathway. Li appears to have moderate effects on downregulation of MAPK & PKC signaling and upregulation of PKA & AMPK signaling, alongside more profound effects on Akt signaling (*Fig. 4F)*. A similar signaling profile (across 3972 phosphoproteins) was observed in BD LR *vs.* BD LR Li, with only 16 differentially phosphorylated proteins linked to the effects of Li (*Fig. 4G, H)*. Of these 16 proteins, 6 were upregulated and 10 downregulated (*Fig. 4G and extended Fig. 4E, F, G)*. Of note, the specific phosphosites (*Fig. 4I)* we link here to Li effect in BD LR patient neurons contain many that are consistent with those implicated in previous BD GWAS (“*GWASdb SNP-Disease Associations” in harmonizome*) and Li phosphopretomic studies (*ref:* https://esbl.nhlbi.nih.gov/Databases/Lithium-Phospho-IMCD/)(Fig. 4I).

Together, our phosphoproteomic results indicate that BD patient neurons exhibit upregulation of the small G-proteins that act as molecular switches to control several cellular processes including: gene expression, cytoskeleton reorganization, vesicular transport, and intracellular signal transduction. Furthermore, the data argue that Li’s beneficial effects in BD LR patient neurons is through downregulation of MAPK and PKC pathways, and upregulation of AMP and Akt pathways.

### Akt activation phenocopies Li effects in BD LR patient neurons, and AMPK activation rescues elevated neural network activity and sodium conductance in BD LNR patient neurons

To test if the pathways we identified by phosphoproteomics can be modulated to mimic the positive effects of Li in BD LR patient neurons, we assessed the neural network activity following inhibition of MAPK (U0126), PKC (GF109203X), and activation of Akt (SC79), AMPK (A769662) signalling over the same time frame as Li application (5-7 days).

Neural network excitability was assayed by imaging activity-dependent somatic Ca^2+^ transients with Fluo4-AM *(Fig. 5A&B and extended Fig. 5A)*. Comparisons between Ctl, BD LR, and BD LNR in non treated and drug conditions are displayed separately for clarity *(Fig. 5A&B)*, but statistical testing is reported from comparison of all groups (*extended Fig. 5A*). Compared to Ctl, BD LR and BD LNR patient neurons exhibited significantly higher Ca^2+^ event frequency *(Fig. 5C)*, as predicted by hyperexcitability (*Fig. 1E)* and spontaneous sEPSC frequency (*extended Fig. 2)* in BD patient neurons. In BD LR patient neurons, Li, Akt, and AMPK activators significantly reduced Ca^2+^ event frequency, whereas no change was observed with PKC or MAPK inhibition *(Fig. 5C)*. This phenocopy suggests Li may act mainly via Akt and AMPK pathways to reduce neuronal activity and achieve its therapeutic effect in BD LR patient neurons. In BD LNR patient neurons, Li had no effect on the neural network activity (*Fig. 5D)*, and MAPK inhibition increased event frequency (*Fig. 5D)*. Interestingly, the activation of AMPK did reduce Ca^2+^ event frequency in BD LNR patient neurons, as was also observed in BL LR patient neurons *(Fig. 5D)*. This suggest targeting AMPK activation could provide an alternative treatment for LR and LNR BD patients.

**Figure 5:**
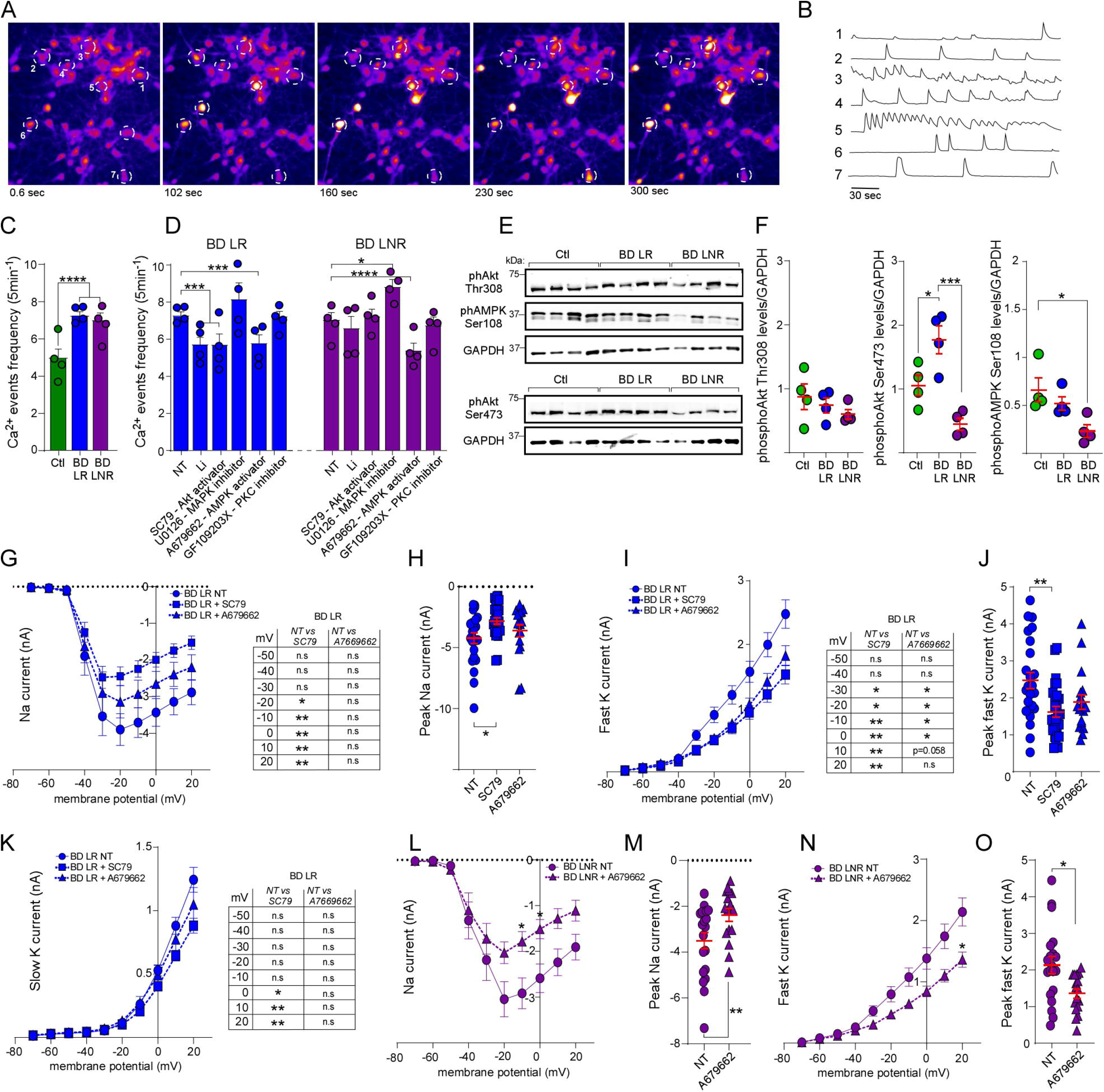
Increased network activity and Na+ & K+ channel hyperactivity in BD neurons is rescued by Akt & AMPK activation. Representative timelapse of Fluo4 Ca^2+^ imaging of human neurons. ***B)*** Representative traces showing Ca^2+^ events (dF/F transients) from individual neurons (ROIs white doted lines in A). ***C)*** Quantification of Ca^2+^ event frequency in Ctl & BD patient neurons in non-treated (NT) condition. ***D)*** Quantification (within group) of NT neurons and those chronically (5 to 7 days) exposed to Li (1.5mM), SC79-akt activator (5uM), U0126-MAPK inhibitor (10uM), A679662-AMPK activator (1uM), or GF109203X-PKC inhibitor (2uM). Data are mean±SEM of 4 lines per group/condition (500 – 800 cells per condition) at 30 to 35 days post-neuronal differentiation. Statistics: Two-way ANOVA with Bonferroni’s multiple comparisons test. *p< 0.05, ***p< 0.001, ****p< 0.0001. ***E-F)*** Representative immunoblot (***E***) and quantification (***F***) of phosphorylated Akt (Thr308) and (Ser473), and AMPKβ (Ser108), normalized to GAPDH. Data shown means ±SEM of 4 different lines per group at ∼30 days post-neuronal differentiation. Statistics: one-way ANOVA with Bonferroni’s multiple comparison post-test ***p< 0.001, *p< 0.05. ***G-O)*** Whole-cell voltage clamp recording of Na (***G, H, L, M***) and K (***I, J, K, N, O***) currents in BD LR & BD LNR patient neurons in non-treated (NT) or chronically applied SC79-akt activator (5uM) or A769662-AMPK activator (1uM). Data are mean±SEM of 20-27 neurons per condition, from 3-4 lines per group at 35-40 days post-neuronal differentiation. Statistics: Two-way ANOVA for ***G***, ***I***, ***K***, ***L***, ***M*** or one way ANOVA for ***H***, ***J***, ***M***, ***O*** with Bonferroni’s multiple comparisons test. **p< 0.01, *p< 0.05.

To validate Akt and AMPK dysregulation in BD patient neurons, relative to Ctl, we quantified phosphorylation residues that report their activation state at Akt Thr308 & Ser473, and AMPK Ser108 by western blot *(Fig. 5E&F)*. No changes were observed in Akt Thr308, whereas Akt Ser473 was hyperphosphorylated in BD LR patient neurons, and hypophosphorylated in BD LNR patient neurons, relative to Ctl *(Fig. 5E, F)*. A differential phosphorylation of Akt Ser473 between BD LR and BD LNR groups may explain the opposite response in BD LR vs BD LNR neurons to Li and Akt activation *(Fig. 5C&D)*. Reduced AMPK Ser108 phosphorylation was found in BD patient neurons but was pronounced and statistically significant only in BD LNR patient neurons, relative to Ctl *(Fig. 5E, F)*. This may explain why BD neurons were responsive to AMPK activation *(Fig. 5D)*, whereas Ctl neurons were not *(extended Fig. 5A)*.

Together, network activity results confirm the predicted hyperactivity in BD patient neurons, and the selective rescue by Li only in BD LR patient neurons. Moreover, the data argue that Li acts via Akt pathway activation in BD LR neurons to trigger its therapeutic effect, and that this can be replicated in BD LR and BD LNR patient neurons by AMPK activation.

Finally, we sought to confirm that activators of Akt (SC79) and AMPK (A679662) have the predicted capacity to mimic Li effects on Na^+^ and K^+^ current in BD neurons. In BD LR neurons, the Akt activator (SC79) significantly reduced Na^+^ and K^+^ currents relative to non-treated neurons (Fig. 5G-J). The AMPK activator (A679662) decreased Na^+^ current and K^+^ in both BD groups (Fig.5G-O) but peak Na+ current reductions were significant only in BD LNR (Fig.5L-O), suggesting that BD LNR neurons are more responsive to AMPK activation, as predicted by pronounced AMPK hypophosphorylation (Fig. 5F). Together the data confirm Akt & AMPK hypoactivity is linked to elevated Na^+^ and K^+^ conductance in BD patient neurons, and that pharmacological activation of Akt & AMPK can reduce neuronal hyperactivity in BD LR & LNR patient neurons.

## DISCUSSION

This study provides mechanistic insights into BD pathophysiology and Li’s mode of action, with direct clinical implications. Building from our observed physiological differences in BD neurons and linking clinical responses to Li with a specific rescue of neural dysfunction, we used transcriptomics and phosphoproteomics to reveal a molecular signature of BD and the specific alterations induced by Li in responsive patient neurons. Focussing on kinase pathways implicated by our ‘omic data, subsequent biochemical and pharmacological data argue that Akt activation may serve as an alternative treatment strategy to Li in BD LR patients, and that AMPK activation may serve as a novel treatment strategy for BD LNR patients.

We confirmed that young (3-to 5-week post-neural differentiation) BD patient neurons exhibit hyperexcitability ^18, 19, 29^ via an increase in APs, and altered Na^+^ channel conductance. We additionally found network / synaptic hyperactivity in BD patient neurons, as evidenced by elevated activity dependent Ca^2+^ bursting and spontaneous synaptic excitatory postsynaptic currents. Action potential, Na^+^ current, and Ca^2+^ bursting hyperactivity in neurons from BD LR patients were all reversed by 7-day Li treatment, but neurons from BD LNR remained hyperactive.

The BD phosphoproteome showed upregulation of small G-protein activity, which was rescued by Li in BD LR patient neurons. Small G-proteins modulate receptors and ion channel activity by influencing both channel opening probability and channel trafficking to the plasma membrane^30^. It is plausible that surface traffic and/or activity of neurotransmitter receptors, ion channels, and ion transporters would be dysregulated in BD LR neurons. Magnesium (Mg^2+^) is an essential cofactor for GTP-binding protein functions and is essential for guanine nucleotide binding and GTP-hydrolysis^31^. Li competes with Mg^2+^ ^32^, and thus could interfere with GTPase binding and activity. There is also known crosstalk between GTPases and Akt signalling pathways^33–35^, which might explain why Akt activity is upregulated in BD LR and downregulated in BD LNR neurons (*Fig. 5E, F*). Further studies could directly assess GTPase activity in BD LR and LNR neurons and subsequent effects on neurotransmitter receptors, ion channel activity, and availability at the plasma membrane.

In contrast to the results here, a recent report^36^ found decreased neuronal excitability human cortical spheroids (hCS) from BD patient hiPSCs after ∼180 days post-differentiation; in which Li treatment increased neuronal activity. The discrepancy with previous published work^18, 19, 29^, could be due to differences in the models’ systems utilized, or reflect specific BD phenotypes at different stage of neuronal development and maturation. Suppose a biphasic neuronal excitability phenotype exists in BD neurons (depending on maturity). In that case, Li can correct it in either direction, depending on the need to reduce or increase activity.

Individuals with BD have 2-fold increased risk of type II diabetes and higher levels of insulin resistance^37, 38^. Recent reports suggested Li exerts its therapeutic effect in BD by acting on insulin signaling^28, 39^. Akt and AMPK kinases are involved in insulin signaling and development of insulin resistance, and we found these to be dysregulated in BD neurons.

Phosphorylation of Akt Ser473 was increased in BD LR patient neurons and decreased in those of BD LNR patients. The differential phosphorylation of Akt Ser473 could explain the differential responses to both Li and Akt activation in BD LR vs LNR patient neurons. We found Li and the Akt activator reduced AP firing, Na^+^, and K^+^ currents, exclusively in the BD LR patient neurons in which they were initially increased. The results predict Akt activation can mimic the therapeutic effect of Li in BD LR patients. In line support of this, a recent report found activated Akt reduced peak Na^+^ current and neuronal excitability in rat cortical neurons^40^.

Elsewhere, inhibition of Akt activity increased Na^+^ current and neuronal excitability in mouse CA1 hippocampal neurons^41^. Notably, Li activates Akt and this activation modulates mood-related behavior in mice^42^. Further, growing evidence^43–46^ shows Akt-related kinases are implicated in schizophrenia and BD; specifically the activity of the Akt pathway is altered in the forebrain of BD and schizophrenia subjects^43^. Together the literature and data here predict Akt activation may be therapeutically useful, but to our knowledge Akt activation is yet to be tested in BD or other neuropsychiatric disorders.

We also found reduced phosphorylation of AMPK at Ser108 in BD patient neurons, with a clearest difference in those from BD LNR patients. It is proposed that reduced AMPK activity contributes to the development of neuropsychiatric disorders and sleep disturbances^47^. Here, an AMPK activator reduced AP hyperactivity, Na^+^ and K^+^ – channel activity in BD LNR patient neurons. AMPK regulates Na^+^ and Ca^2+^ channels, and the Na/K pump^48, 49^ responsible for maintaining neural membrane polarity and thus excitability. Activation of AMPK reduces excitability via inhibition of Na^+^ channels, regulates K^+^ channels leading to hyperpolarization, and dampens L-type Ca2+ channel currents^50, 48, 49^. Here AMPK activation reduced neural activity in neurons from BD LR and LNR patients; although reductions of Na^+^ and K^+^ current were clearer in BD LNR neurons suggesting they are more responsive to AMPK activation. A plausible explanation is our observation that AMPK activity was lower in BD LNR than BD LR neurons at base, making cellular effects of AMPK activation easier to affect. An alternative explanation, supported by a previous report^29^, is that hyperexcitability in BD LNR neurons is related to a reduction of Wnt/b-catenin signaling, and consequent downregulation of b-catenin/TCF/LEF1 transcriptional activity. This study showed that Li had no effect on b-catenin in BD LNR cells, maintaining low b-catenin/TCF/LEF1 transcriptional activity, therefore, Li could not reduce the hyperexcitability in BD LNR. Interestingly, b-catenin and its transcriptional activity are stabilized by AMPK phosphorylation of b-catenin at Ser 552^51^; crosstalk between Wnt/b-catenin and AMPK signaling could explain why AMPK activation dampened neural hyperactivity in BD LNR neurons.

A recent study found metformin (an AMPK activator) improved clinical outcomes in BD patients^52^. Specifically, BD patients who responded well to metformin reported improved mood disorder symptoms and decreased insulin resistance. In that study, metformin reduced depressive symptoms; while this provides support for the use of AMPK activation in BD, further studies are needed to confirm the results.

In summary, by utilizing hiPSC technology and personalized neuronal disease modeling, we found neuronal hyperactivity in BD patient neurons that was corrected by Akt and AMPK activators. We propose these as targets for alternative treatment strategies in BD for further testing. Future studies with larger samples and especially those including female patients, will aim to corroborate the results here, and guide preclinical and clinical studies. This work also provides the framework for a personalized drug screening platform to accelerate development of other alternative therapeutic strategies for BD. Finally, the capacity to distinguish between BD LR and BD LNR patient’s neuronal response to Li (or other treatments) and predict a beneficial clinical response for individuals. As such this study advances our capacity to realize personalized medicine for BD to close the gap between diagnosis and therapeutic intervention, minimize the interim suffering, and drastically reduce the risk of suicide.

## METHODS

### Ethics statement

All cell lines and protocols in the present study were used in accordance with guidelines approved by the institutional human ethics committee guidelines and the Nova Scotia Health Authority Research Ethics Board (REB # 1020604).

### iPSC line generation and neuronal differentiation

All the iPSCs were reprogrammed from blood samples of consenting individuals. The iPSCs lines from healthy individual (SBP009, SBP011, SBP012), BD LR patients (SBP005, 007, 010) and BD LNR (SBP001, 002, 004) were reprogrammed from lymphocytes as described previously^19^. The other iPSC lines were reprogrammed from peripheral blood mononuclear cells (PBMCs) using CytoTune iPS Sendai reprogramming kit (Thermo Fisher Scientific) according to the manufacture’s protocol with the adequate quality control criteria for iPSC validation.

iPSCs were cultured on a Matrigel (Corning) coated plate with mTeSR1 medium (STEMCELL Technologies) and neural progenitor cells (NPCs) induction was performed according to Stem cell Technologies protocol with STEMdiffTM SMADi Neural Induction kit (STEMCELL Technologies). On Day0, a single-cell suspension of iPSC was generated using Gentle Cell Dissociation reagent and 10,000 cells per microwell was seeded in an ultralow attachment 96-well plate to generate Embyoid bodies (EBs), cultured in STEMdiff™ Neural Induction Medium + SMADi + 10 μM Y-27632. From Day1 to Day4, a daily partial (3/4) medium change was performed using STEMdiff™ Neural Induction Medium + SMADi. On Day5, EBs were harvested using a wild-board 1ml serological pipettes and 40μm strainer and transferred to a single well of a 6-well plate coated with Poly-L-ornithine hydrobromide (PLO) and laminin before. A daily full medium change was performed from Day6 to Day11. After the neural induction efficiency was determined higher than 75%, Neural rosettes are manually selected using STEMdiff™ Neural Rosette Selection Reagent on Day12 and replated onto a single well of PLO/Laminin coated 6-well plate. With continuous daily full medium change, selected rosette-containing clusters attached, and NPC outgrowths formed a monolayer between the clusters.

Neural Progenitor Cells (NPCs) were ready for passage 1 when cultures are approximately 80 – 90% confluent (typically at Day17 to Day19). NPCs were maintained using STEMdiff™ Neural Progenitor Medium and ready to differentiate after two passages with Cell Gentle Dissociate Reagent. For the final differentiation into Hippocampal neurons, NPCs were digested with Accutase and then differentiated onto PLO/laminin coated plates/coverslips with final differentiation media which is BrainPhys media (STEMCELL Technologies) supplemented with Glutamax (Fisher), N2, B27, 200nM ascorbic acid (STEMCELL Technologies), 500 μg/ml cyclic AMP (SIGMA), 20 ng/ml brain-derived neurotrophic factor (BDNF, GIBCO), 10 ng/ml Wnt3a (R&D Systems), and 1 μg/ml laminin for 2 weeks; with a medium change of 3 times per week. After 2 weeks of differentiation, final differentiation media is replaced by STEMdiffTM forebrain Neuron Maturation kit until neurons are used. Maturation media were changed twice a week.

### Immunocytochemistry and confocal imaging

iPSC-derived neurons were fixed in phosphate-buffered saline (PBS) containing 3.7% formaldehyde and 5% sucrose for 1 h at room temperature (RT), then in NH4Cl (50mM) for 10 min. Neurons were then permeabilized for 20 min in PBS containing 0.1% Triton X-100 and 10% goat serum (GS) at RT and immunostained with a guinea-pig polyclonal anti-vglut1 (1/4000; Millipore AB5905), a mouse monoclonal anti-BIII tubulin (1/1000; Sigma T8660), a mouse anti-Sox2 (1/500 Abcam ab79351), a rabbit anti-Nestin (1/600 Sigma N5413) and DAPI (1/5000 Sigma D9542), antibodies in PBS containing 0.05% Triton X-100 and 5% GS. Cells were washed three times in PBS and incubated with the appropriate secondary antibodies (1/1000) conjugated to Alexa 555 or Alexa488 and mounted with Prolong (ThermoFisher P36930) until confocal examination.

The images (1024L×L1024) were acquired with a ×20 lens (numerical aperture NA 1.4) on an SP8 confocal microscope (Leica Microsystems). Z-series of 6 images of randomly selected field were compressed in 2D using the max projection function in image j (FIJI).

### Electrophysiological recordings and analyses

All electrophysiological signals were acquired using Multiclamp 700B amplifier digitized at 10kHz and pClamp10 software. Data were analyzed in Clampfit10 (Molecular Devices).

#### External and internal solutions

iPSC-derived neurons on coverslips were transferred to a recording chamber in a recording buffer containing (in mM): 167 NaCl, 10 D-glucose, 10 HEPES, 2.4 KCl, 1 MgCl_2_, and 2 CaCl_2_ (300-310 mOsm, pH adjusted to 7.4 with NaOH) supplemented by SC79 (5uM, Tocris 4635) or A769662 (1uM, Sigma SML2578) depending on the experimental condition. Whole cell patch clamp experiments were carried out at 25–27 °C. For all recordings, pipettes were filled with a potassium-based solution containing (in mM): 145 K-gluconate, 3 NaCl, 1 MgCl_2_, 1 EGTA, 0.3 CaCl_2_, 2 Na-ATP, 0.3 Na-GTP, 0.2 cAMP and 10 HEPES (290 mOsm, pH adjusted to 7.3 with KOH). Patch pipettes displayed a resistance of 4– 7LMΩ.

#### Synaptic events recordings

sEPSCs and sIPSCs were recorded in voltage clamp mode, holding cells at -55 and 0 mV respectively. Synaptic events were sampled for 5min and then exported to Clampfit 10.7 software with which the amplitude and frequency of synaptic events were quantified (threshold 10pA); all events were checked by eye and monophasic events were used for amplitude and decay kinetics, while others were suppressed but included in frequency counts as in these studies^53, 54^. Tolerance for series resistance (Rs) was <30 MΩ and uncompensated; ΔRs tolerance cut-off was <20%.

#### Sodium and potassium currents

Sodium and potassium currents were acquired in voltage-clamp mode. Sodium channel currents are reported as inward peak currents and potassium channel currents as outward currents during series of voltage steps of 10mV from -70mV to 20mV. Currents were directly normalized by the cell capacitance during recordings in pClamp10 software.

#### Neuronal excitability

For assessing neuronal excitability, action potential (AP) firing was recorded in current-clamp mode in response to incremental, hyperpolarizing followed by depolarizing current injections of 1s duration (3pA increment of 15 steps). The number of AP firing was plotted to the corresponding current steps using Clampfit 10.7 software. In current-clamp mode, the resting membrane potential of all cells was adjusted to ∼ -65mV by injection of a small negative current if needed.

### RNA extraction and RNA sequencing

RNA sequencing was performed on bulk cultures of neurons at ∼ 30 days post-differentiation treated or not with Li for 7 days in 5 Ctl, 5 BD LR and 4 BD LNR patient lines (28 samples). Total RNA was extracted from ∼3x10^6^ cells for each condition using miRNeasy kit (Qiagen, USA) according to the manufacturer’s instructions. RNA was resuspended in RNAse-free water. The RNA concentration was measured on the Synergy H4 microplate reader. RNA was sent to Genome Quebec center for sequencing. Library preparation was done using the TruSeq Stranded Total RNA Kit (Ilumina) with Ribo-Zero depletion. Sequencing was done on the NovaSeq 6000 at 150bp paired end reads with a total of 113M reads (∼17Gb/sample).

### Differential expression analysis and Pathway enrichment

The count alignment and quantification were performed with STAR (v2.7.3a) and RSEM (v1.3.1). The reference genome used was GRCh38.p13. The quantified raw counts were transformed to log2CPM with Limma package (version 3.56.0). Genes with at least one count per million (CPM) in more than 25% samples were retained. Samples having connectivity with other samples were retained (Z-scored standard deviation <3.5). Limma was used for differential expression analysis. We separately compared gene expression levels between Ctl and BD lines, as well as BD LR and BD LNR -/+ Li. Significant genes (FDR<0.05, -log10(pvalue)>2, logFC>1) were further enriched with GO and KEGG pathways (clusterProfiler R package version 4.8.1).

### Phosphoenrichment and Phosphoproteomic analysis

#### Protein lysis and digestion

∼30 days post-differentiation iPSC-derived neurons were collected in 50 ul of Tris-buffer saline (TBS) solution supplemented with protease inhibitor cocktail (Sigma P8340) and phosphatase inhibitors (PhosSTOP, Sigma 4906845001). ∼3x10^6^ cells per condition (∼1.2mg of proteins). For each sample (nL=L4 per group), 50LuL of cell pellets were lysed, reduced, and alkylated in lysis buffer (8 M Urea, 2 mM DTT, 100 uM Orthovanadate, supplemented with 10 mM iodoacetamide (IAA) after half an hour). The samples were then diluted in 50 mM ammonium bicarbonate to a Urea concentration of 1 M. Proteins were digested overnight at 25L°C with trypsin (Promega) with an enzyme/substrate ratio of 1:250. After digestion, samples were supplemented with 2 volumes of acetonitrile (ACN) and precipitate (DNA/RNA) was eliminated after a 10 min centrifugation in an Eppendorf centrifuge at 14,000Lrpm.

#### Ti02 phosphoenrichment

The supernatant was supplemented with 6% Trifluroacetic acid (TFA) and transferred to Eppendorf tubes containing 10 μL of 5 μm TiO2 beads (Canadian Biosciences) and incubated for 30 min on a rotating wheel. The supernatant was aspirated and the TiO2 beads were washed twice with 50% ACN/0.5% TFA in 200LmM NaCl and once with 50% ACN/0.1% formic acid (FA). Phosphopeptides were eluted with 10% ammonia in 50% ACN. Samples were dried down in a speed vac, resuspended in 20 μL of 0.1% FA in water and stored frozen, if not processed immediately.

#### Mass spectrometry: RP-nanoLC-MS/MS

Mass spectrometry data were acquired using an UHPLC Easy nLC 1000 (Thermo Scientific) coupled to an Orbitrap Q Exactive HF mass spectrometer (Thermo Scientific). 50% of the phosphopeptide enriched samples were first trapped (Acclaim PepMap 100 C18, 3 μm, 2Lcm) before being separated on an analytical column (Acclaim, C18 2 μm, 25Lcm). Trapping was performed in solvent A (0.1% FA in water), and the gradient was as follows: 2-20 % solvent B (0.1% FA in 90%ACN. 10% water) in 90Lmin, 20-38 % in 60Lmin, 38-90% in 10Lmin, maintained at 90% for 10Lmin, then, back to 0% solvent B in 10Lmin. The mass spectrometer was operated in data-dependent mode. Full-scan MS spectra from m/z 375–1500 were acquired at a resolution of 120,000 at m/z 400 after accumulation to a target value of 5 × 106. Up to 25 most intense precursor ions were selected for fragmentation. HCD fragmentation was performed at normalized collision energy of 35% after the accumulation to a target value of 1 × 105. MS/MS was acquired at a resolution of 30,000. A dynamic exclusion was set at 6Lseconds.

#### Bioinformatics and Databases for Protein Identification (BioID) and analysis

All MS/MS raw data files were converted into peak lists using Mascot Peak Distiller (Matrix Science). Mgf files were searched against the curated human proteome database (UniProt: Homo sapiens) using the Mascot search engine (Matrix Science) with the following parameters: enzyme: trypsin. Missed cleavages: 1. #C13:1. Fixed modifications: C-acetamidation. Variable modifications: oxM and phospho-T and phospho-S. Precursor mass accuracy: 6 ppm. MSMS mass accuracy: 50 mmu. Instrument type: ICRFT. Mascot files were imported into Scaffold v5 (Proteome Software Inc), and a second search engine (X!Tandem) was requested to search the data again, after which the combined data was validated through the trans proteomic pipeline. Data were visualized as TSCs in scaffold v5 software, counting only the phosphopetides identified with protein threshold of 20%, Min # peptides of 1 and peptide threshold of 95% and batched in categories according to patient and experimental parameters. (Ctl, BD LR, BD LNR -/+ Li in all groups). For the quantitative method, we used Total spectra with minimum value of 1 and normalization. When the comparison is >2 groups, one-way ANOVA test was used, otherwise t-test to identify the differential phosphoproteins with p< 0.05.

### Somatic calcium imaging

iPSC-derived neurons were washed with Krebs HEPES Buffer and incubated with 1 μm Fluo 4-AM (Thermofisher F14201) in Krebs HEPES Buffer for 20 min at room temperature. Excess dye was removed by washing twice with Krebs HEPES Buffer, and cells were incubated for an additional 20 min to equilibrate the intracellular dye concentration and allow de-esterification.

Then, cells were in Krebs HEPES buffer supplemented with the different drugs used for the Ca^2+^ experiment described in Fig. 4C. Live-cell imaging was performed by excitation of fluo4 at 488nm laser and measured at 520nm at 25-26 °C. A Time-lapse image sequences of 500 frames were acquired at ∼2 Hz with a region of 512pixels × 512 pixels using a Andor digital camera (Andor solis). Images were subsequently processed using EZCalcium^55^ in Matlab software (MATLAB R2022b, MathWorks) to automatically identify the region of interest (ROI) which is the soma of the cells. Then, CaSiAn^55^ software implemented in Java was used to quantify the large Ca^2+^ events frequency in a semi-automated manner.

### Immunoblot

Cells were homogenized in lysis buffer (10 mM Tris–HCl pH7.5, 10 mM EDTA, 150 mM NaCl, 1% Triton X100, 0.1% SDS) in the presence of a mammalian protease inhibitor cocktail (Sigma, P8340) and phosphoSTOP (Sigma 4906845001) to protect proteins from dephosphorylation. Protein extracts (30μg) were resolved by SDS–PAGE, transferred onto PVDF membrane (Millipore IPFL00010), immunoblotted with the indicated concentration of primary antibodies: rabbit monoclonal anti-phospho-Akt (Thr308) (1/1000, Cell Signaling #4056), rabbit monoclonal anti-phospho-Akt (Ser473) (1/1000; Cell Signaling #4060); rabbit monoclonal anti-phospho-AMPKβ1 (Ser108) (1/1000; Cell Signaling #23021). Standard GAPDH loading controls were included using a mouse monoclonal anti-GAPDH antibody (1/2000, ThermoFisher MAB5-15738). Then membrane was revealed using the appropriate LI-COR fluophore-conjugated secondary antibodies. Images were acquired on a LI-COR Odyssey Infrared image system. Fluorescence intensity values for each protein of interest were normalized to GAPDH signal from the same gel. Full-size blots for cropped gels can be found in extended data.

### Drug treatment

The therapeutic range of Li treatment is between 0.75-1.5mM. In this study, neurons were treated chronically with **∼**1.5mM Li (Sigma L9650) for 7 days. Akt activator SC-79 (Tocris 4635) has been used in the literature^56–58^ between 5 and 10uM. We chose to use low concentration (5uM) but chronic treatment as Li treatment. A769662 (Sigma SML2578) is a potent activator of AMPK with half maximal effective concentration (EC50) of ∼1uM^59^. Thus, we used this concentration for our experiments. Based on the literature and several assays used previously, we used MAPK inhibitor U0126 (Tocris 1144) at 10uM^60, 61^ and PKC inhibitor GF109203X (Tocris 0741) at 2uM^62^. All the concentrations were chosen to activate or inhibit ∼30 to 50% the kinase activity during 5 to 7 days.

### Data manipulation and statistical analyses

Statistical analyses were performed using GraphPad Prism9 software (GraphPad software, Inc). All data are expressed as mean ± standard error of the mean (s.e.m.). Normality for all groups was verified using the Shapiro-Wilk tests and p<0.05 was considered significant. All statistical tests used are described in the figure legend.

## Data availability

All relevant data are in the figures and supplementary figures. Any raw data and materials that can be shared will be released via a material transfer agreement. All transcriptomic data from this study are deposited in the GEO repository (GSE236482) and proteomic data will be deposited in the ProteomeXchange Consortium.

## ACKNOWLEDGMENTS

We gratefully acknowledge the financial supports from ERA PerMed grant to MA and G.A.R, Bell Let’s Talk-Brain Canada Mental Research Program to AK, AM, MA, and G.A.R, Brain & Behavior Research Foundation Young investigator award 2022 to AK. We also thank the microscopy platform of the Montreal Neurological Institute (MNI) and the Proteomics & Molecular Analysis Platform, RIMUHC.

## AUTHOR CONTRIBUTIONS

MA provided the patient samples. AK and YL developed the protocol for differentiation from IPSC to Neural Precursors then to forebrain neurons. YL grew and differentiated all the lines. AK performed the staining and imaging of neurons, analyzed by AK and LS. AK, MAb and AKa performed the electrophysiological recordings. AK and MAb performed the Ca^2+^ imaging experiments. AK collected the RNA samples, CJ and QH analyzed the RNA sequencing. AK collected the protein samples and KD performed phosphoenrichment, AK and KD analyzed the phosphoproteomic data. AK collected protein samples and YC and AP performed the Western Blot. CEC and NF participated in methods development. AK and A.J.M contributed to study design, curation and development, and data interpretation. AK, G.A.R and A.J.M provided the overall supervision and funding. AK wrote the original draft edited by LS and A.J.M and revised by all authors.

