## supplementary figures for "Molecular signatures of hyperexcitability and lithium responsiveness in bipolar disorder patient neurons provide alternative therapeutic strategies"

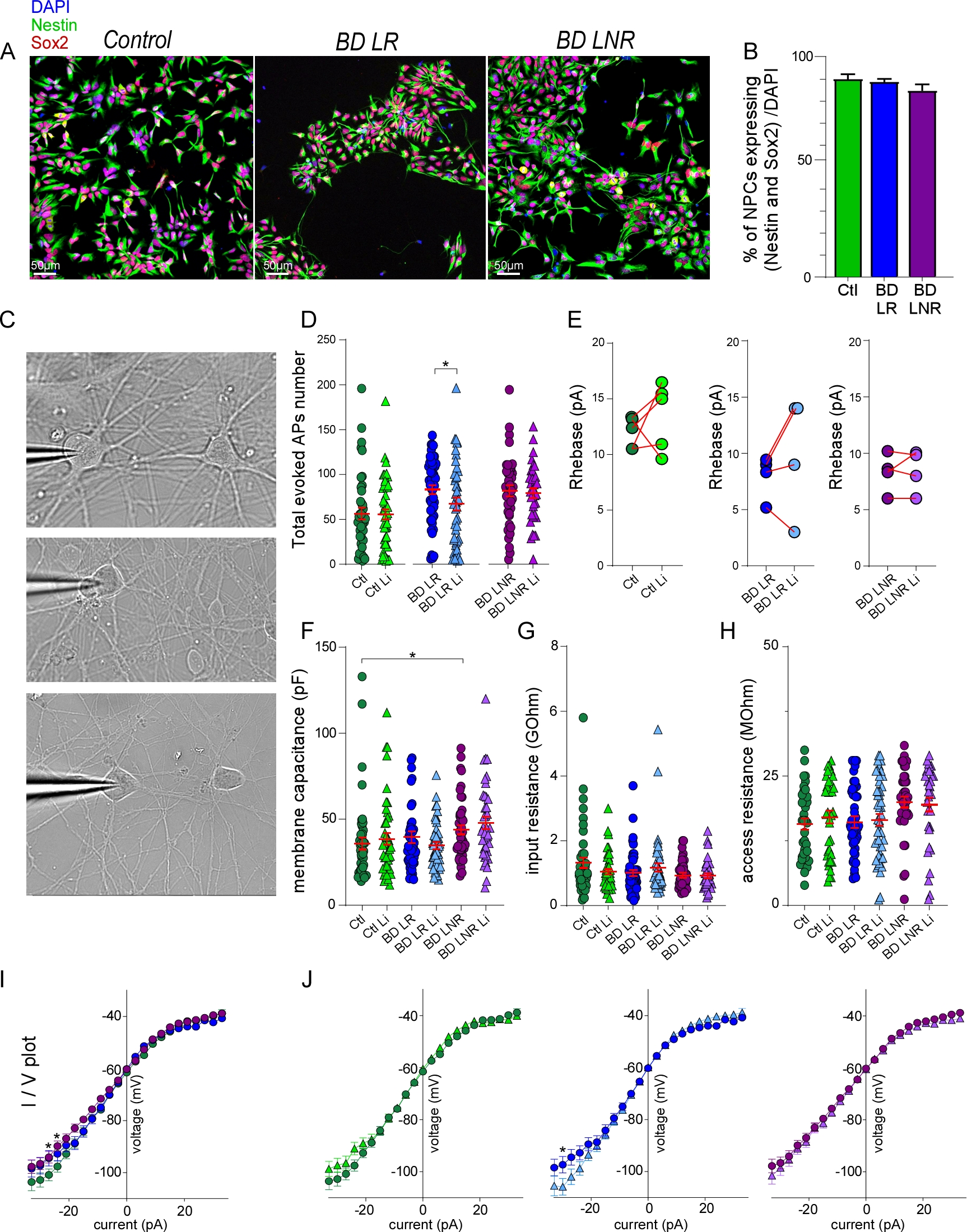

**Extended data figure 1**: **Neural precursors characterization and measurements of membrane properties.**

***A)*** Nestin and Sox2 markers were used to confirm an efficient production of Neural Precursor Cells (NPCs) for each group. ***B)*** Quantification of well-differentiated NPCs represented as a percentage of NPCs expressing both Nestin and Sox2 (~90%). Data shown are the mean ±SEM of ~14 to 19 images per condition from 4-5 different lines per group. Statistics: non significance was determined by ANOVA with Bonferroni’s multiple comparisons test. ***C)*** Representative excitatory neurons recorded from, with large soma (> 30um) exhibiting proper dendritic extensions. ***D)*** Scatter plots show quantification of total evoked action potentials of neurons from each group -/+ Li for 7 days in response to a 1 second depolarizing of 3pA current injection steps from 0 to 33 pA. Data shown are the mean ±SEM of ~50 neurons per condition from 4-5 different lines per group/condition and statistical significance determined by non-parametric Mann-Whitney test for the comparison of -/+ Li conditions in each group. *p< 0.05. ***E)*** Rheobase represented by the mean of each line -/+ Li for each group showing more in details how each line responds to the treatment in each group. Data shown are the means of 4-5 different lines per group/condition. ***F-H)*** Scatter plots show quantification of membrane properties prior current injection steps. ***I-J)*** Voltage-current curves for each group -/+ Li (1.5mM) for 7 days with a membrane resting potential at ~-60mV. Data are the mean ± SEM of ~50 neurons per condition from 4-5 different lines per group and statistical comparison were one-way ANOVA for F-H and two-way ANOVA for I-J with Sidak’s multiple comparisons test. *p< 0.05

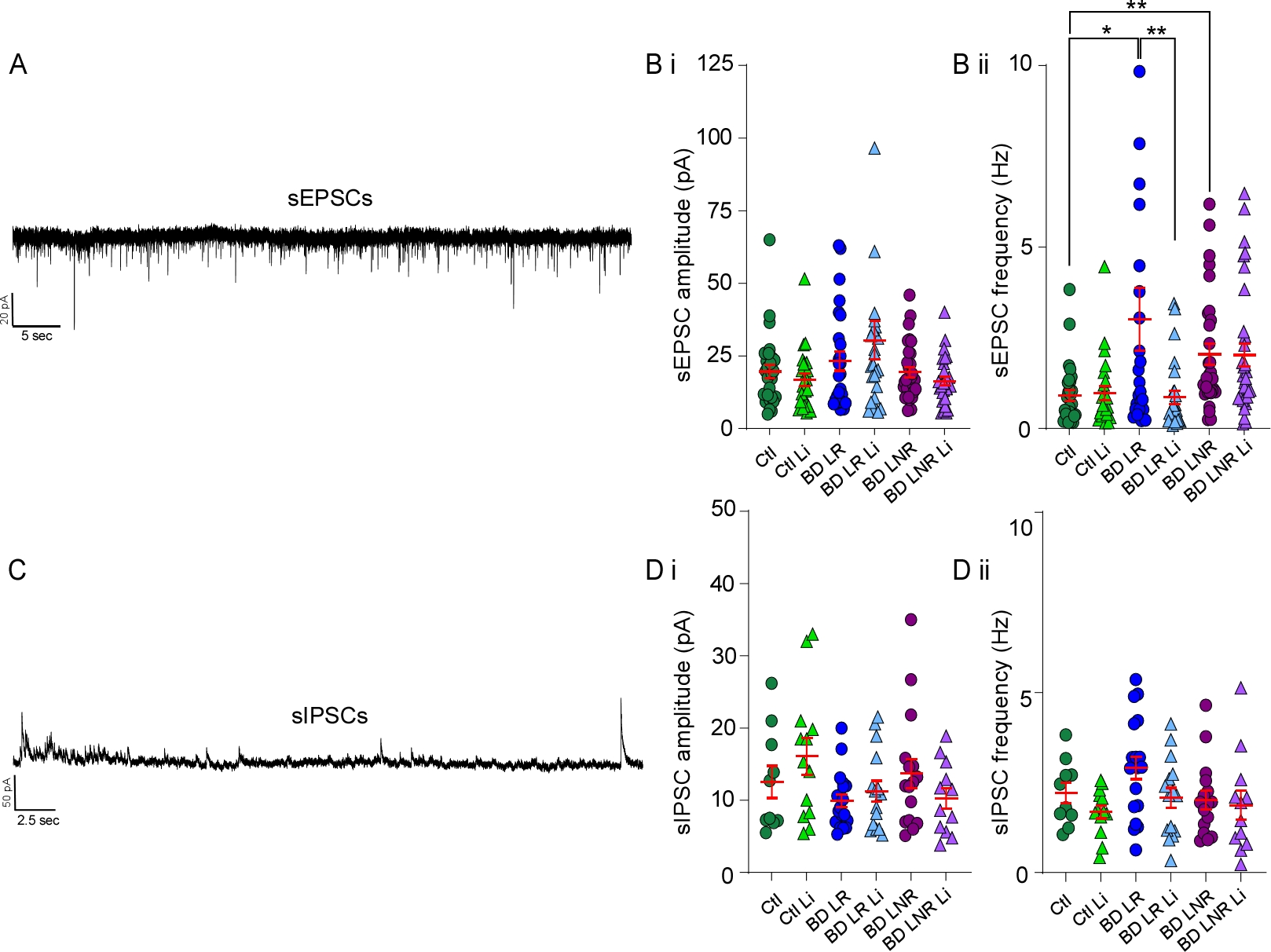

**Extended data figure 2**: **Increased spontaneous excitatory postsynaptic current frequency in BD neurons that is selectively rescued by Li in BD LR patient lines.**

A) Representative sample traces of sEPSCs from human neurons. Scatter plot show quantification of amplitude in Bi and frequency in Bii of sEPSCs of ~30 neurons per condition from 4-5 different lines per group. C) Representative sample traces of sIPSCs from human neurons. Scatter plot show quantification of amplitude in Di and frequency in Dii of sIPSCs of ~15 neurons per condition from 4-5 different lines per group. Data shown in B and D are the mean ± SEM and statistical significance determined by non-parametric Kruskal-Wallis’s test with Dunn’s multiple comparison post-test. *p< 0.05, **p< 0.01.

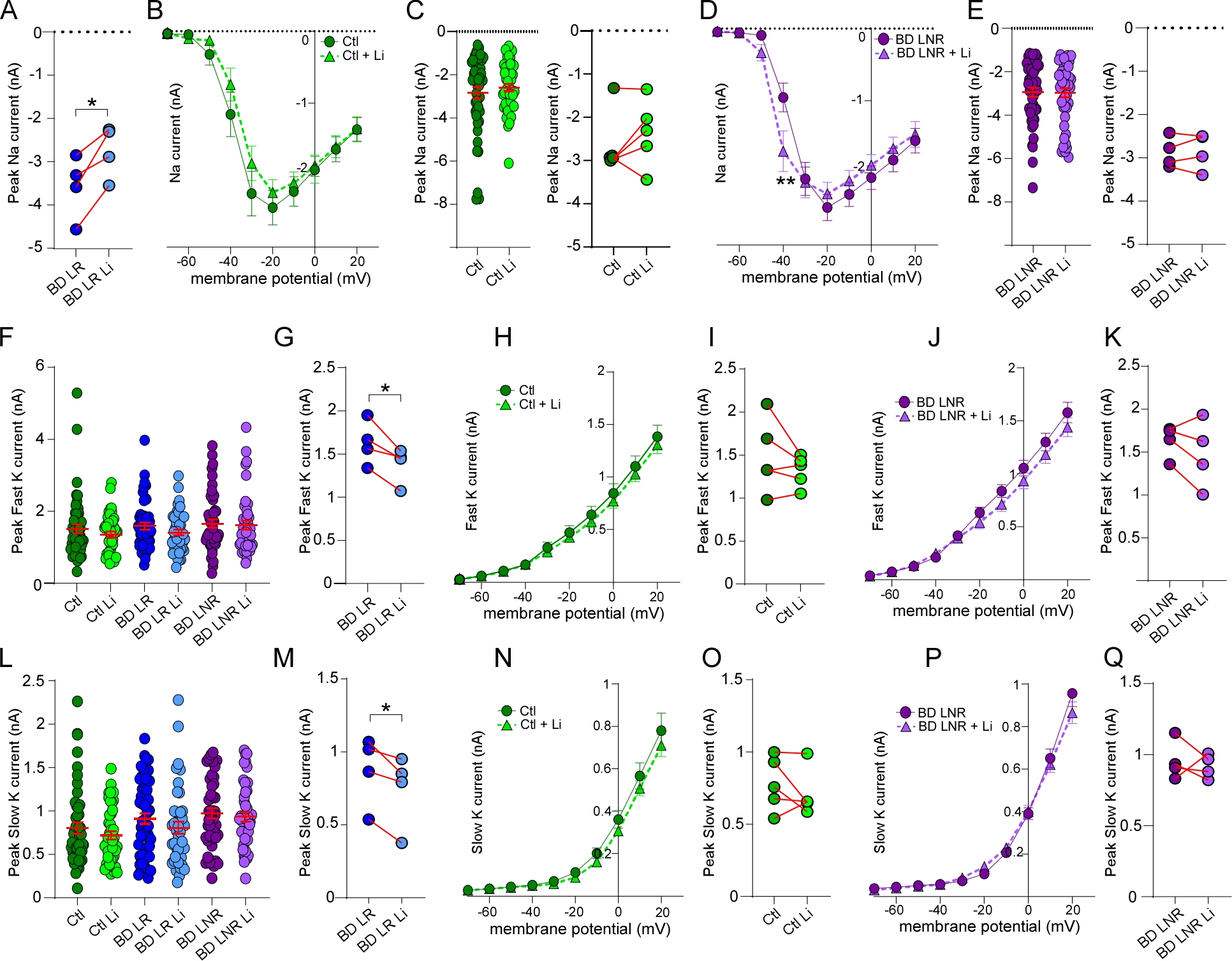

**Extended data figure 3**: **Measurement of voltage-dependent sodium and potassium currents in control and BD LNR -/+ Li.**

***A)*** Peak of sodium current represented by the mean of each line -/+ Li from 4 BD LR patients showing that all the BD LR lines respond to the Li treatment by reducing their sodium current. Voltage-dependence of sodium current in ***B*** and ***D)*** and peak sodium current in ***C*** and ***E)*** in iPSC-derived neurons from control and BD LNR neurons at 28 to 32 days post-neuronal differentiation treated or not chronically with Li (1.5mM) for 7 days. ~50 neurons per condition from 4-5 different lines per group/condition. The means per line represented in ***C*** and ***E*** -/+ Li from 5 Ctl and 4 BD LNR patients show how Li affects sodium current in each line of these groups. Statistic: Two-way ANOVA for ***B***, ***D*** and one-way ANOVA for ***C***, ***E*** with Bonferroni’s multiple comparisons test. Paired t test was used with the means per lines in ***A***, ***C*** and ***E***. *p< 0.05.

Quantification of peak of fast and slow potassium currents in ***F*** and ***L)*** -/+ Li conditions in all groups. Quantification of voltage-dependence of fast and slow potassium currents in control (***H-N***) and BD LNR patients (***J-P***). Data shown are means ±SEM of ~50 neurons per condition from 4-5 different lines per group at 28 to 32 days post-neuronal differentiation. The means per line represented in ***G***, ***I***, ***K*** and ***M***, ***O***, ***Q*** -/+ Li from 5 Ctl, 4 BD LR and 4 BD LNR patients show how Li affects fast or slow potassium currents in each line of these groups. Statistic: Two-way ANOVA for ***H***, ***J***, ***N***, ***P*** and one-way ANOVA for ***F***, ***L*** with Bonferroni’s multiple comparisons test. Paired t test was used with the means per lines in ***G***, ***I***, ***K***, ***M***, ***O*** and ***Q***. *p< 0.05.

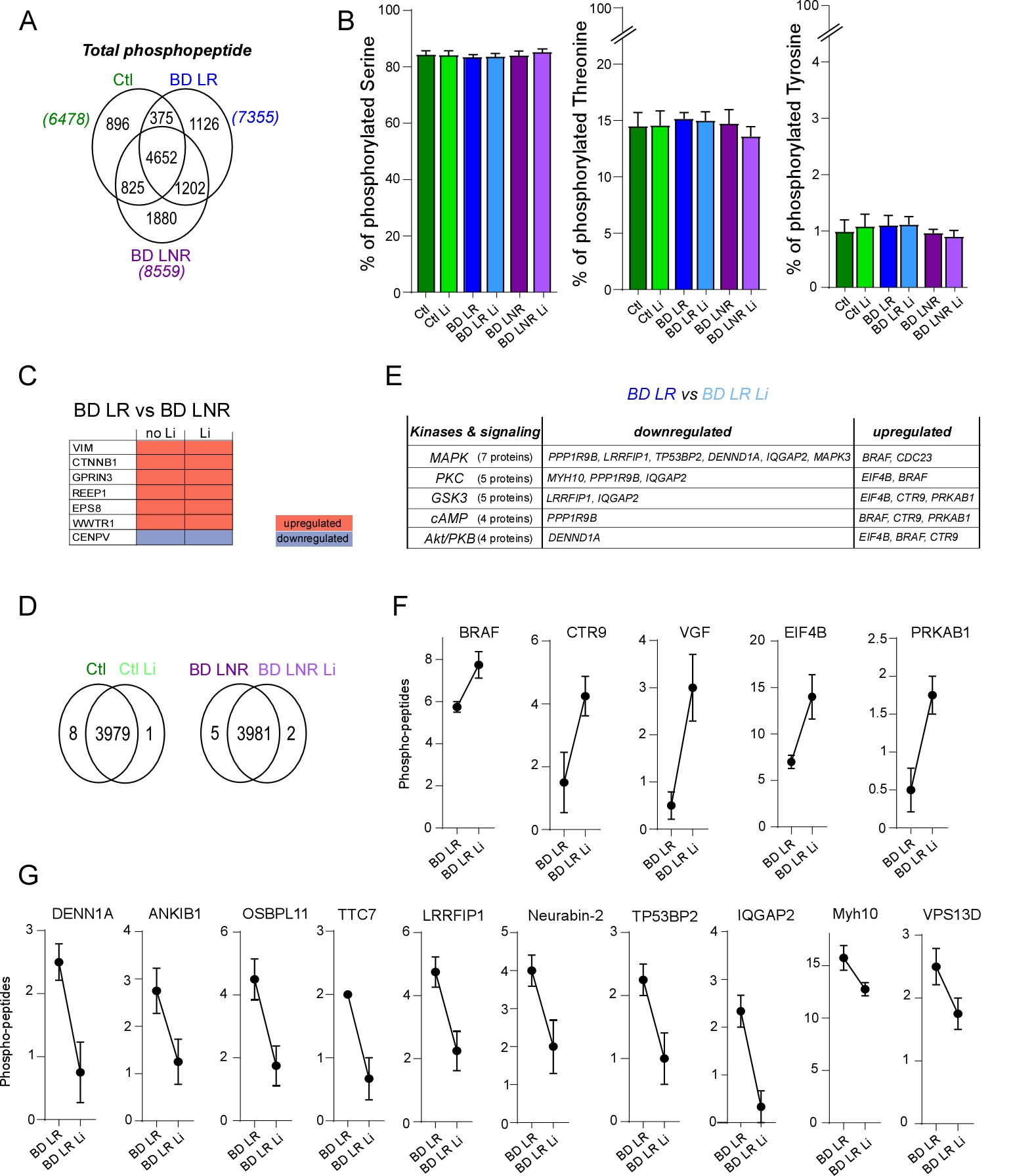

**Extended data figure 4: Comparison of identified phosphopeptides in all groups -/+ Li treatment.**

***A)*** Venn diagram shows the number of phosphopeptide identified in each group on average from 4 different lines per group. ***B)*** Bar graphs show the percentage of identified phospho-Serine, phospho-threonine, and phospho-tyrosine sites in all groups -/+ Li treatment. Data are the mean ± SEM of 4 different lines per group/condition and statistical non-significance was determined by ANOVA with Bonferroni’s multiple comparisons test. ***C)*** Heatmap of differential phosphoproteins in BD LR vs BD LNR between non- and Li-treated conditions showing an overlap of 7 phosphoproteins between -/+ Li conditions, which means not affected by Li. ***D)*** Venn diagram shows the number of differential phosphoproteins in Ctl and BD LNR in non-treated and Li conditions. ***E)*** Table shows the number of identified substrates phosphorylated by MAPK, PKC, GSK3, cAMP and Akt signaling and whether each signaling is up or downregulated by Li treatment in BD LR group. ***F-G)*** The significant upregulated (in ***F***) or downregulated (in ***G***) phosphoproteins by Li in BD LR lines by comparing the number of phosphopeptides identified. Data shown are means ±SEM of 4 different lines per group/condition with statistical significance of p< 0.05 determined by paired t-test.

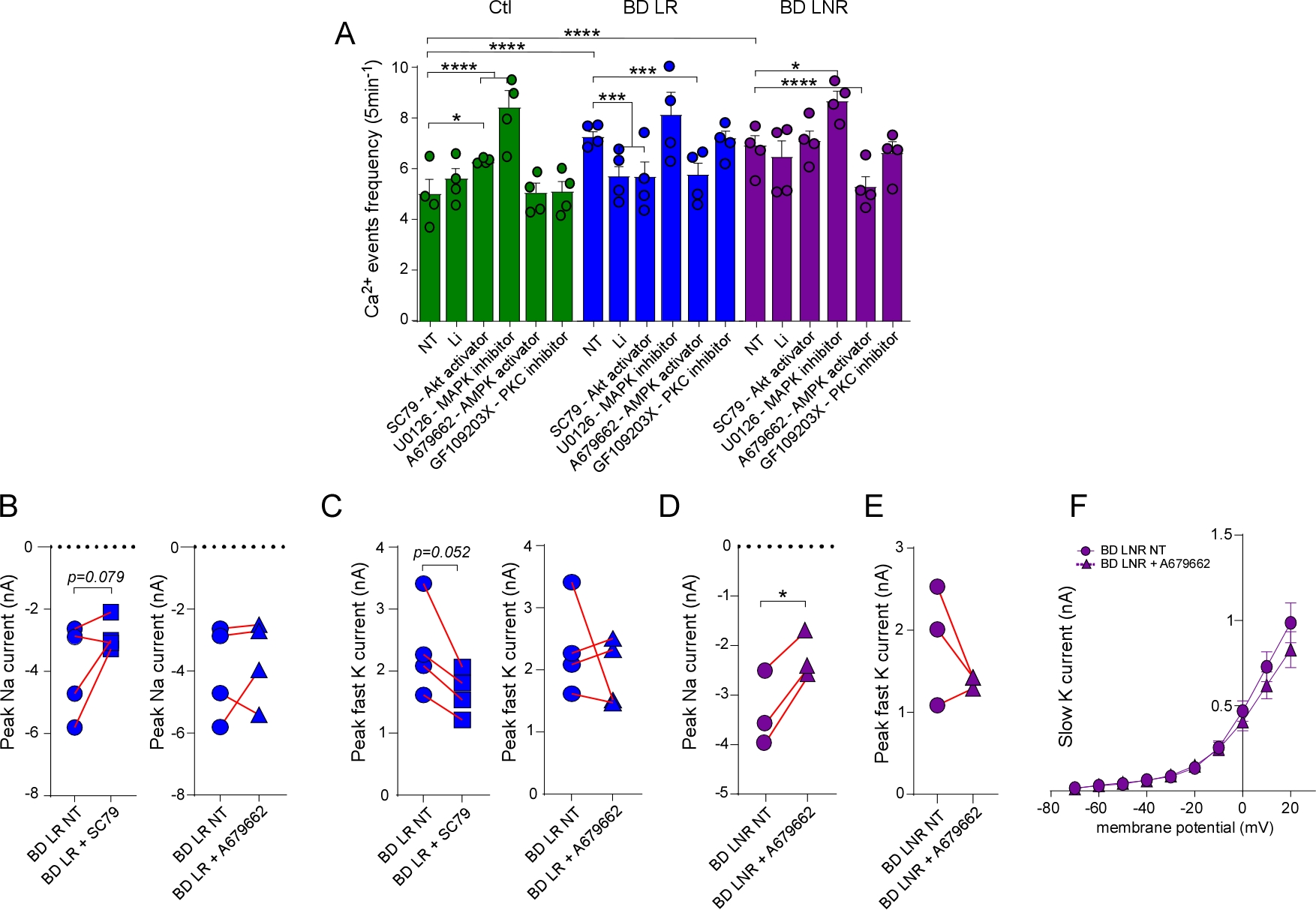

**Extended data figure 5: Decreased voltage-dependent sodium currents in BD LR and LNR with SC79 and A679662 treatment.**

***A)*** Bar graphs show quantification of Ca^2+^ event frequencies in Ctl, BD LR and BD LNR neurons in non-treated (NT) condition or treated chronically (5 to 7 days) by Li (1.5mM), SC79-akt activator (5uM), U0126-MAPK inhibitor (10uM), A679662-AMPK activator (1uM), GF109203X-PKC inhibitor (2uM). Data shown are means ±SEM of 4 different lines per group/condition at 30-35 days post-neuronal differentiation with 500 to 800 cells analyzed per condition. Statistics: two-way ANOVA with Bonferroni’s multiple comparisons test. *p< 0.05, ***p< 0.001, **** p<0.0001. ***B-C)*** Peak of sodium or fast potassium currents represented by the mean of each line -/+ SC79 or A679662 from 4 BD LR patients showing how BD LR neurons respond to the chronic treatments by reducing their sodium and potassium currents. ***D-E)*** Peak of sodium or fast potassium currents represented by the mean of each line -/+ A679662 from 3 BD LNR patients showing how BD LNR neurons respond to chronic A679662-AMPK activator treatment by reducing their sodium and potassium currents. No changes in Voltage-dependence of slow potassium current -/+ A679662 treatment in BD LNR group. Data shown are means ±SEM of 20 to 27 neurons per condition from 3-4 different lines per group at 35-40 days post-neuronal differentiation. Statistics: Two-way ANOVA for ***F*** or paired t-test for ***B***, ***C***, ***D*** and ***E***. *p< 0.05.
